# Replication-coupled search positions MutH for strand incision in DNA mismatch repair

**DOI:** 10.64898/2026.07.21.739762

**Authors:** Amy N Moores, Nehir Banaz, Tendai Makamure, Stephan Uphoff

**Affiliations:** Dorothy Crowfoot Hodgkin Building, Department of Biochemistry, University of Oxford, South Parks Road, Oxford OX1 3QU

## Abstract

DNA mismatch repair (MMR) safeguards genome stability by correcting replication errors, yet how its components coordinate as an efficient pathway in living cells remains unclear. In *Escherichia coli*, MutS and MutL detect mismatches and must rapidly activate MutH to incise the nascent DNA strand at distantly located GATC sites. Here, we used live-cell single-molecule tracking to resolve the target-search dynamics of MMR proteins *in vivo*. We find that MutH binds DNA at hemimethylated GATC sites independently of mismatch recognition and without MutS or MutL. The autonomous recruitment of MutH is facilitated by a switch from fast diffusion to a slow-search mode near DNA replication forks. This guidance allows MutH to transiently occupy potential incision sites in the wake of replication forks, awaiting activation by MutS–MutL. The parallelised search mechanism of MMR proteins facilitates rapid and targeted repair of replication errors in cells.

## Introduction

All organisms from bacteria to humans rely on molecular machines to ensure accurate replication of their genomes. DNA mismatch repair (MMR) plays a central role in the maintenance of genome stability by correcting misincorporated nucleotides before they become fixed as mutations^1,2^. The proteins that perform MMR face the remarkable challenge of detecting rare mismatches among millions of correctly paired bases in the genome. In the absence of MMR, mutation rates increase by 100- to 1000-fold^3,4^, and loss of MMR is a major driver of disease, contributing to cancer development^5,6^ and drug resistance^7^ in humans, and antibiotic resistance ^8,9^ and host adaptation^10,11^ in pathogens. Accordingly, the MMR pathway has been studied intensively for decades. Genetic, biochemical, and structural approaches have identified the components of mismatch repair and elucidated their key mechanisms^2^. More recently, single-molecule approaches have provided crucial dynamic insights *in vitro*^12,13^. However, these approaches have not yet explained how MMR proteins operate together inside living cells to achieve rapid and accurate repair.

Each step in the pathway depends on precisely regulated protein–DNA and protein–protein interactions controlled by ATP-dependent conformational changes. Although this pathway was originally characterised in *Escherichia coli*, its core principles are conserved throughout evolution. In *E. coli*, MMR is initiated by the coordinated actions of MutS, MutL, and MutH^14^. A MutS dimer first recognises a mismatch and recruits a MutL dimer, which subsequently activates the MutH endonuclease^15–17^. MutL alters the conformation of MutH to introduce a nick at a GATC sequence on the newly synthesised strand of the DNA duplex. Following incision, a cascade of additional repair enzymes unwind, degrade, and resynthesise the stretch of DNA from the GATC cut site to replace the misincorporated base.

Crucially, the MMR enzymes need to distinguish a mismatch from the vast excess of correctly paired DNA. MMR not only avoids nicking and degrading undamaged DNA but also fixes >99% of mismatches correctly^4^. A defining feature of MMR is its ability to distinguish the newly synthesised strand containing the error from the parental template. In *E. coli* and related γ-proteobacteria, this strand discrimination is mediated by DNA methylation. GATC sequences are methylated by DNA adenine methylase (Dam) within seconds to minutes after replication at most sites^18,19^, creating a transient time window during which GATC sites are hemimethylated in the wake of the replication fork. MutH exploits this asymmetry by selectively incising the unmethylated daughter strand. Because Dam-mediated methylation occurs rapidly, the MMR machinery must detect mismatches and activate MutH before the strand-discrimination signal is lost. Failure to do so is the main cause of persistent mismatches that become permanent mutations in the subsequent round of DNA replication^20^.

In addition to this kinetic constraint, the MMR pathway faces a fundamental spatial challenge. GATC sites are often located hundreds to thousands of base pairs away from a mismatch, on either side of the mismatch. How MutS, MutL, and MutH coordinate across these distances to position MutH at the correct incision site remains a central longstanding question^21^. Several contrasting mechanisms have been proposed, including one-dimensional sliding of MutS and MutL along DNA^22^, trapping and assembly of multimeric MutL along DNA^23^, or DNA looping to bring distant sites into proximity^24,25^. Central to these models is the question of whether MutH binds GATC sites independently awaiting activation, or is recruited to the mismatch region by the other mismatch repair proteins. Although MutH preferentially incises the GATC site nearest the mismatch^26,27^, this trend does not distinguish whether reduced incision at distal sites reflects inefficient delivery or activation of MutH over longer distances. *In vitro* studies have provided evidence supporting aspects of each model, but no consensus has emerged. Importantly, such experiments cannot fully capture the native chromosomal organisation or the dynamic cellular context in which DNA replication, methylation, and repair occur simultaneously.

To fill this knowledge gap, we employed live-cell single-molecule tracking^28–30^ to directly monitor the intracellular activities of MutS, MutL, and MutH during mismatch repair. This approach enables the observation of individual repair proteins as they move through the cell and bind their targets. Our experiments focussed on establishing how MutH is recruited and positioned to execute strand incision, the key committed step of the MMR pathway.

## Results

### Live-cell single-molecule tracking uncovers DNA-binding and target-search populations of MMR enzymes

To enable live-cell single-molecule tracking, we generated *E. coli* strains expressing HaloTag fusions of the mismatch repair proteins MutS, MutL, or MutH from their native chromosomal loci. To verify that the fusion proteins retained mismatch repair activity, we quantified spontaneous chromosomal mutation frequencies using the emergence of rifampicin-resistant colonies as a readout (Fig. S.1.). All HaloTag fusion strains displayed mutation frequencies indistinguishable from the wild type progenitor. In contrast, deletion of *mutS*, *mutL*, or *mutH* resulted in ∼100-fold elevated mutation frequencies, consistent with hypermutator phenotypes of MMR-deficient cells^4^. These data demonstrate that the HaloTag fusions preserve full mismatch repair functionality.

We labelled Halo-tagged proteins in exponentially growing cells using the bright cell-permeable synthetic fluorophore Tetramethylrhodamine (TMR), which undergoes stochastic photoswitching between fluorescent and dark states^31^. This enabled short-lived, high-frame-rate observations (∼30-70 frames s⁻¹ over ∼15 frames) of many individual MutS, MutL, and MutH molecules moving within live cells. Localisations in successive frames were linked into trajectories to reconstruct protein movements, and diffusion coefficients were subsequently extracted from the mean-squared displacement of each trajectory^32^. The resulting diffusion coefficient distributions reveal the target search dynamics of MutS, MutL, and MutH for mismatch repair sites (Fig 1).

**Fig. 1:**
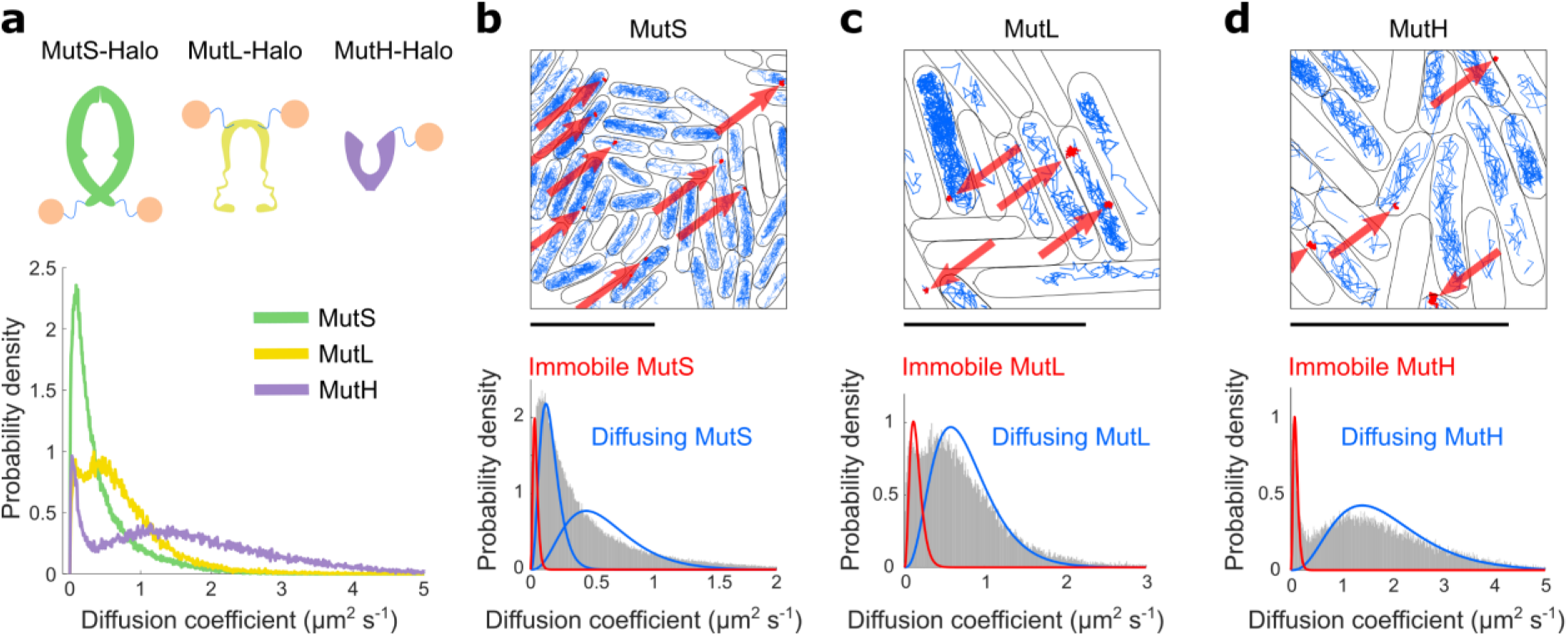
MMR proteins exhibit distinct binding and search dynamics in vivo. **a,** Schematic of HaloTag fusions to MutS, MutL, MutH. Distribution of diffusion coefficients of MutS-Halo-TMR, MutL-Halo-TMR, MutH-Halo-TMR in live *E. coli* cells. Data are pooled from 3 independent experiments for MutS and MutL, and 6 experiments for MutH. **b, c, d,** Top: Representative cells (segmented outlines, black) with single-molecule tracks showing diffusing (blue) and immobile molecules (red) for MutS-Halo-TMR, MutL-Halo-TMR, MutH-Halo-TMR. Arrows indicate binding events. Scale bars, 5 µm. Bottom: Distribution of diffusion coefficients with fit curves for immobile and diffusing states of MutS-Halo-TMR, MutL-Halo-TMR, MutH-Halo-TMR. Data are pooled from 3 independent experiments for MutS and MutL, and 6 experiments for MutH.

For all three proteins, we observed a mixture of immobile and diffusing molecules in cells (Fig 1a-d). The fraction of molecules belonging to each population was estimated by fitting the distributions of single-molecule diffusion coefficients with a two-state model comprising an immobile and a diffusing population (Fig 1b-d). For MutS, the diffusing population contained a slow and a fast species of molecules, as previously characterised^30^. Among the mobile populations, MutS exhibited the lowest diffusion coefficient, with mobility pattern matching previous reports in *B. subtilis*^29^ and *E. coli*^28,30^. MutL showed intermediate mobility and MutH diffused most rapidly (Fig 1d). These differences are consistent with the relative molecular weights of the proteins (97 kDa, 70 kDa, and 25 kDa for MutS, MutL, and MutH, respectively^15,33^), as well as their known oligomeric states: MutS and MutL predominantly function as dimers (or tetramers)^1,2^, whereas MutH acts as a monomer^17^.

Directly imaging active mismatch repair events at the single-molecule level in live cells is challenging, as mismatches are rare, occurring at a rate of ∼0.1 per cell cycle^4,34,35^, and repair is completed within minutes^20,34^. Consistent with this, only a small fraction of trajectories for each protein were classified as immobile under basal conditions: 10.4% (±0.4%) for MutS, 19.8% (±5.2%) for MutL, and 11.3% (±1.2%) for MutH (errors: ± s.e.m.). While immobile trajectories may correspond to proteins bound at mismatch repair sites under these conditions, they could also reflect non-specific DNA interactions during target search^36^ or association with other cellular components such as the replication machinery^29,37^. To determine which of these interactions are related to active mismatch repair, we next perturbed the pathway and examined how the binding behaviour of MutS, MutL, and MutH responds to increased mismatch formation.

### MutH recruitment occurs independently of MutS and MutL

To increase the number of molecules actively engaged in mismatch repair, we treated cells with the DNA-damaging agent methyl methanesulfonate (MMS). MMS generates a range of DNA lesions, including O⁶-methylguanine, which leads to G–T mismatches that are recognised by the MMR pathway^35^. Following MMS treatment, the fraction of immobile MutS and MutL molecules increased significantly (Fig 2a-b), consistent with increased mismatch binding by MutS and subsequent recruitment of MutL to the site. In contrast, neither the diffusion coefficients nor the immobile fraction of MutH molecules were affected by MMS treatment (Fig 2c). These observations indicate that MutH binding is independent of mismatch presence and suggest that loading of MutS and MutL at mismatches does not directly trigger recruitment of MutH.

**Fig. 2.**
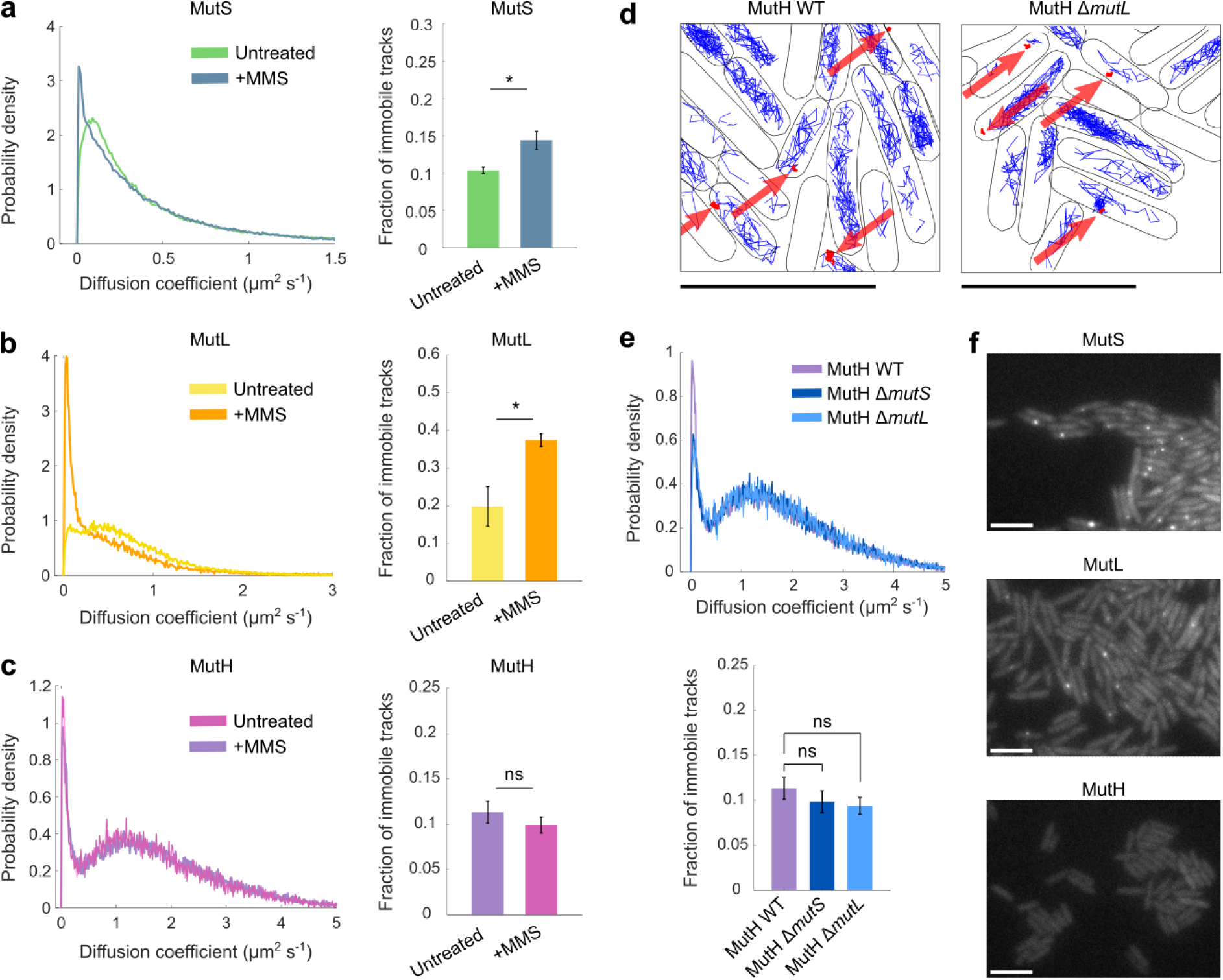
MutH binds irrespective of mismatch recognition and independently of MutS and MutL. **a, b, c** Distribution of diffusion coefficients (left) and fraction of tracks classified as immobile (right) for MutS-Halo-TMR, MutL-Halo-TMR, MutH-Halo-TMR in untreated cells and in MMS-treated cells. Distributions are pooled and bar plots show mean ± s.e.m from 3 independent experiments for all conditions, except MutH Untreated condition which is comprised of 6 independent experiments (*: P<0.05, ns: P>0.05). **d,** Representative cells (segmented outlines, black) with single-molecule tracks showing diffusing (blue) and immobile molecules (red) for MutH-Halo-TMR in wild-type cells and in Δ*mutL* cells. Arrows indicate binding events. Scale bars, 5 µm. **e,** Distribution of diffusion coefficients (top) and fraction of tracks classified as immobile (bottom) for MutH-Halo-TMR in wild-type, Δ*mutS*, Δ*mutL* cells. Distributions are pooled and bar plots show mean ± s.e.m from 3 independent experiments for Δ*mutS* and Δ*mutL* conditions and 6 independent repeats for MutH WT (ns: P>0.05). **f,** Representative fluorescence snapshots of MMS-treated cells expressing MutS-mYPet, MutL-mYPet, MutH-mYPet. Identical greyscale applied to all three images. Scale bars, 5 µm.

To substantiate this theory, we additionally performed single-molecule tracking of MutH in *E. coli* cells where MutS and MutL are absent. If mismatch recognition by MutS and MutL were required for MutH recruitment, deletion of these genes would be expected to reduce MutH binding. However, MutH immobile trajectories were retained under deletion of either *mutS* or *mutL* (Fig. 2d), and no effect was observed on the overall fraction of immobile MutH-Halo molecules (Fig. 2e). Together, these results indicate that the MMR machinery does not assemble through a simple linear hierarchical recruitment of MutS, MutL, and then MutH. Rather, MutH associates with DNA independently of MutS-MutL, and the tripartite activation complex is formed in parallel from pre-bound components.

This conclusion is also supported by snapshot imaging of mismatch repair enzymes fused to the fluorescent protein mYPet (Fig 2f). In MMS-treated cells, MutS-mYPet and MutL-mYPet formed bright, discrete foci, indicating the accumulation of multiple enzyme copies at or near mismatch sites, consistent with previous observations^23,35^. In contrast, MutH-mYPet fluorescence was uniformly distributed throughout the cytoplasm in all cells, and no MutH foci were observed under any conditions. These data argue against models in which MutH associates stably with long-lived stationary or sliding protein complexes during mismatch repair.

### MutH binding depends on active DNA replication

Our data suggest that MutH binds DNA independently of mismatches and of the other MMR proteins. A plausible hypothesis is that MutH associates with newly synthesised DNA, where transiently hemimethylated GATC sites are generated following replication and provide the substrate for strand incision. In this model, MutH would be pre-positioned on the chromosome prior to mismatch recognition, poised for activation by MutL. To test if MutH binding depends on active replication, we halted DNA replication by treating *E. coli* cells with two bacteriostatic antibiotics: chloramphenicol, which inhibits translation, and rifampicin, which inhibits transcription. Although their mechanisms of action differ, the inhibition of protein synthesis by both antibiotics ultimately prevents replication initiation^38^, and thereby reduces the fraction of cells with ongoing DNA replication (Fig S.2.). We performed single-molecule tracking of MutH-Halo following treatment with chloramphenicol or rifampicin for 1 hour. In both conditions, the fraction of immobile MutH-Halo trajectories was significantly reduced in non-replicating cells (Fig 3a-b), showing that MutH binding indeed depends on ongoing DNA replication.

**Fig. 3.**
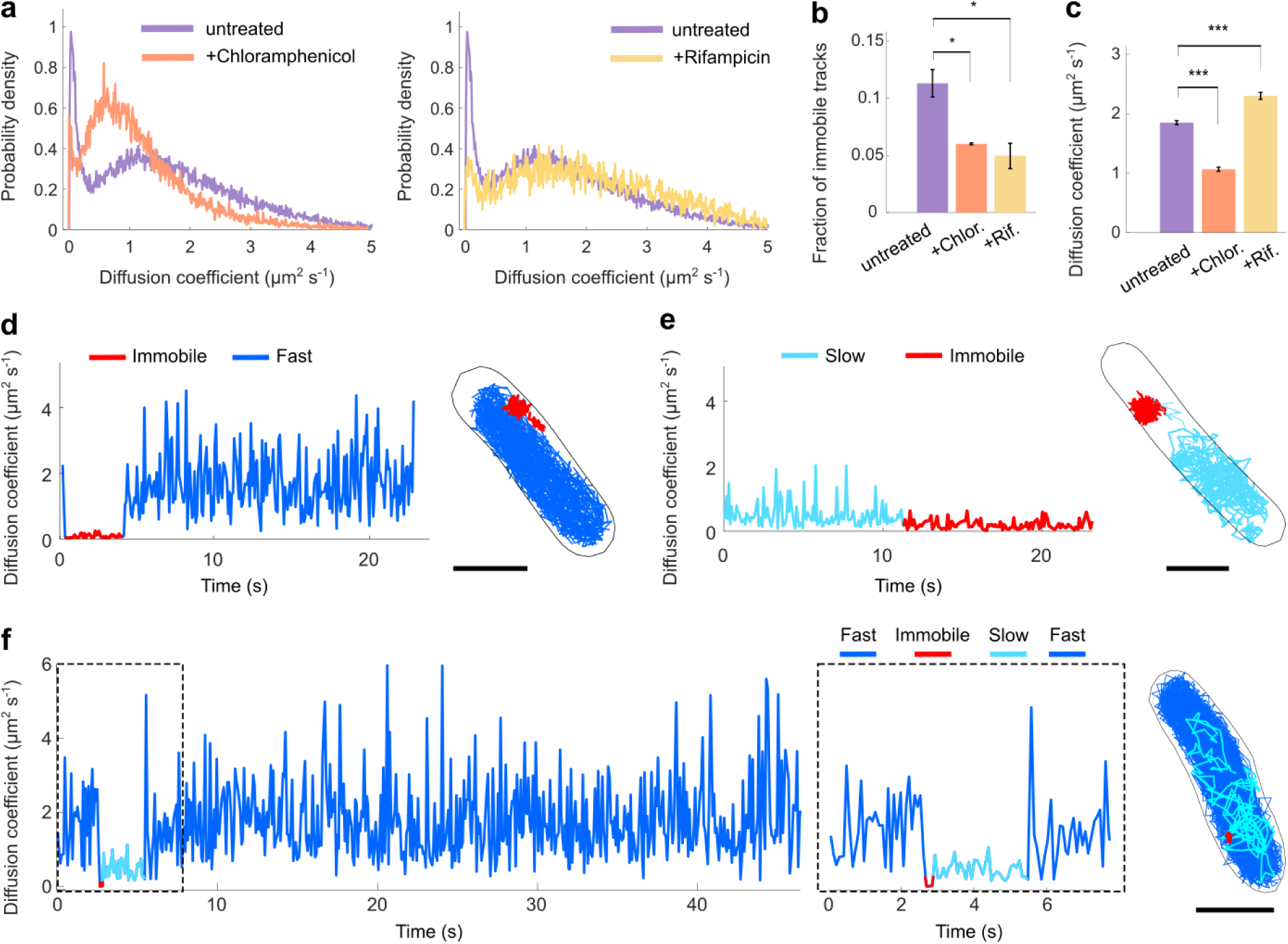
Replication-dependent switching between fast, slow and immobile states of MutH. **a,** Distribution of diffusion coefficients for MutH-Halo-TMR in untreated cells and in cells treated with chloramphenicol (left) or rifampicin (right). Data are pooled from 3 independent experiments for chloramphenicol and rifampicin treated cells, and from 6 independent repeats for the untreated condition. **b,** Fraction of MutH-Halo-TMR tracks classified as immobile in untreated cells and in cells treated with chloramphenicol or rifampicin. Bar plots show mean ± s.e.m across independent experiments (*: P<0.05). **c,** Mean diffusion coefficient of diffusing MutH-Halo-TMR molecules in untreated cells and in cells treated with chloramphenicol or rifampicin. Bar plots show mean ± s.e.m across independent experiments (***: P<0.0005). **d, e, f** Representative time-traces of diffusion coefficients and corresponding tracks of single MutH-Halo-JFX650 molecules showing transitions between fast diffusion (dark blue), slow diffusion (light blue) and immobile (red) states. Scale bars, 1 µm. Dashed box shows magnified part of the time-trace in f.

The antibiotic perturbations also informed about how MutH searches for binding sites, as we observed marked changes in the mobility of diffusing MutH molecules (Fig 3c). In chloramphenicol-treated cells, MutH diffusion slowed substantially (mean D = 1.07 µm^2^s^-1^) relative to untreated cells (mean D = 1.85 µm^2^s^-1^). In contrast, MutH molecules diffused more rapidly in rifampicin-treated cells (mean D = 2.30 µm^2^s^-1^). These opposing changes reflect the contrasting effects of chloramphenicol and rifampicin on chromosome architecture, which condense and expand the nucleoid, respectively^39^. To test the influence of chromosome organisation on MutH target search without inhibiting DNA replication, we expanded the chromosome by deleting the nucleoid-associated protein HU^40^. We observed the predicted increase in MutH mobility and no change in the fraction of immobile tracks (Fig. S.3.).

In contrast, deletion of *mutS* or *mutL* had no detectable effect on MutH diffusion (Fig. 2e). If MutH frequently formed complexes with either protein during target search, an increase in mobility would be expected in their absence. Together, these results indicate that MutH target search is governed primarily by interactions with the chromosome rather than with other MMR proteins. The sensitivity of MutH mobility to DNA density suggests that this search involves frequent non-specific DNA interactions, similar to the behaviour of other DNA-binding proteins^36^.

### Prolonged single-molecule tracking reveals two distinct target-search modes of MutH

We next sought to visualise the trajectories of MutH during its search for and binding to target sites. To extend the observation time of individual molecules, we labelled MutH-Halo with the photostable HaloTag ligand JFX650, enabling continuous tracking of single molecules for tens of seconds to several minutes^30^. These extended trajectories captured complete cycles of target search, DNA binding, and dissociation. Individual MutH binding events had an average duration of 3.3 s (±0.97 s s.e.m., Fig. S4a), indicating that MutH transiently occupies potential incision sites while awaiting activation by MutS–MutL.

Unexpectedly, prolonged observation with JFX650 revealed that MutH adopts two distinct modes of target search. In addition to the immobile state, MutH molecules exhibited two different mobile states: a fast-diffusing state (∼1.7 µm² s⁻¹) and a slow-diffusing state (∼0.5 µm² s⁻¹) (Fig. 3d-f). While the presence of an immobile and a mobile population was already apparent in shorter tracks acquired using TMR labelling, the existence of two distinct mobile states became evident only in long-lived trajectories enabled by JFX650. Individual MutH molecules persisted in either the fast- or slow-diffusing state over extended periods, with discrete transitions between the two modes observable within single trajectories. The slow-diffusing state contributes negligibly to the quantification of the immobile population, validating our earlier two-state classification (Fig. S4b). Both fast- and slow-diffusing states persisted in a Δ*mutL* background (Fig. S4c), indicating that these search modes are intrinsic to MutH and independent of MutL or prior mismatch recognition.

Molecules in the fast-diffusing state typically explored the full cell volume, whereas slow-diffusing molecules appeared spatially confined to the central region occupied by the nucleoid. We therefore propose that the slow-diffusing state reflects a DNA-scanning mode in which MutH remains in close proximity to the chromosome, facilitating efficient recognition of hemi-methylated GATC sites. This interpretation is consistent with structures of MutH^17^, which show that its two domains can pivot relative to each other via a flexible linker, potentially enabling multiple modes of DNA interaction. In contrast to MutS and MutL, which undergo ordered ATP-driven conformational cycles during mismatch recognition and signalling^41–43^, MutH transitions between immobile, fast-, and slow-diffusing states without a defined sequence (Fig. 3d-f). This stochastic switching is consistent with MutH acting independently during target search, without energy-driven coordination of its conformational states.

### MutH switches target search mode and binds near replication forks

To place the two MutH target-search modes and binding events into the spatial context of a replicating chromosome, we labelled DNA replication loci in the same cells. For this purpose, we used a fluorescent fusion of the β-sliding clamp, YPet-DnaN^44^. Sliding clamps form characteristic foci that trail replication forks, as tens of DnaN molecules remain bound at completed Okazaki fragments^45^. We first acquired snapshots of YPet-DnaN foci and subsequently performed single-molecule tracking of MutH-Halo-JFX650 in the same cells. This approach allowed us to relate transitions between MutH mobility states to the locations of newly synthesised DNA, where hemi-methylated GATC sites are expected to be enriched.

Indeed, MutH binding events occurred in the vicinity of DnaN foci (Fig 4a-b). To quantify this spatial association, we measured the distances between MutH binding sites and the nearest YPet-DnaN focus. We compared this distribution to a randomised control in which MutH binding sites were reassigned to random locations within the same segmented cell area. MutH binding events clustered close to DnaN foci, with a mean separation of 0.33 µm (± 0.06 µm s.e.m.), significantly closer than for random localisation (mean separation 0.67 µm ± 0.08 µm s.e.m.). In addition to co-localised binding, we found that transitions of MutH from fast diffusion to the slow-search mode also occurred preferentially near DnaN foci (Fig 4c-d), with a mean separation of 0.32 µm (± 0.04 µm s.e.m.), compared to 0.72 µm (± 0.13 µm s.e.m.) for randomly localised transitions.

**Fig. 4.**
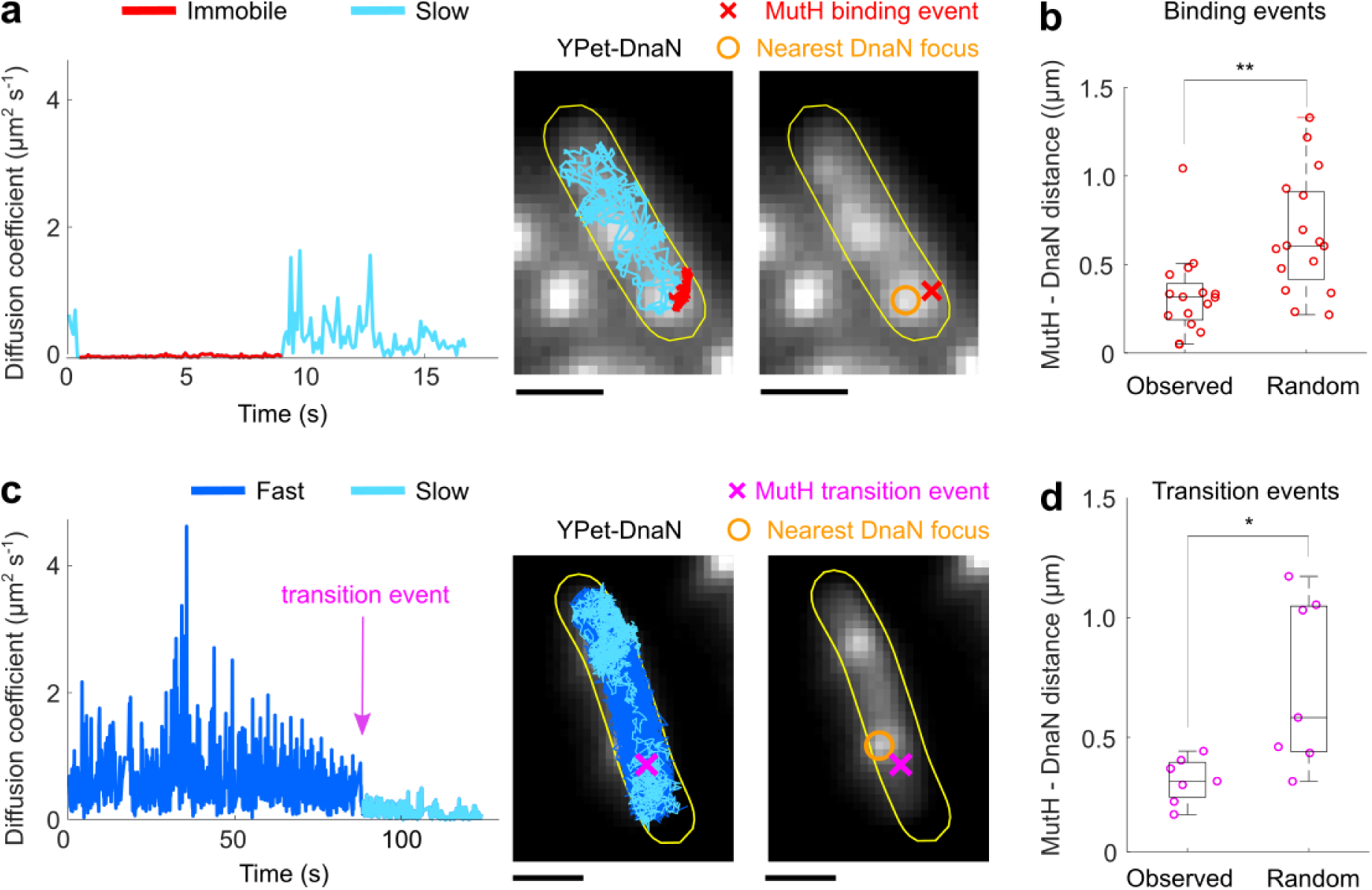
MutH switches between fast and slow search modes and binds near DNA replication forks. **a,** Left: Representative time-trace of diffusion coefficients and corresponding track of a single MutH-Halo-JFX650 molecule showing a binding event (red). Right: Corresponding fluorescence snapshot of YPet-DnaN in the same cell. Localisations of the MutH binding event (red cross) and the nearest DnaN focus (orange circle) are shown. Scale bars, 1 µm. **b,** Box plots of the distances between MutH-Halo-JFX650 binding events and the nearest YPet-DnaN focus per cell. Observed are experimentally measured distances. Random are distances when MutH binding sites were reassigned to random locations within the segmented cell area. ** P<0.005. **c,** Left: Representative time-trace of diffusion coefficients and corresponding track of a single MutH-Halo-JFX650 molecule showing a transition from fast diffusion to slow diffusion. Right: Corresponding fluorescence snapshot of YPet-DnaN in the same cell. Localisations of the MutH fast-to-slow transition event (pink cross) and the nearest DnaN focus (orange circle) are shown. Scale bars, 1 µm. **d,** Box plots of the distances between MutH-Halo-JFX650 fast-to-slow transition events and the nearest YPet-DnaN focus per cell. Observed are experimentally measured distances. Random are distances when MutH fast-slow-transition sites were reassigned to random locations within the segmented cell area. * P<0.05.

### MutH binding is directed by hemimethylated GATC sites

Our data support a model in which MutH changes its search mode to scan and bind newly-replicated DNA near active replication forks. We asked whether this behaviour is specified by hemimethylated GATC sequences. Alternatively, MutH could be recruited via interactions with replisome components, as has been reported for MutS and MutL in multiple species^29,37^. To directly test whether MutH localisation is governed by the availability and methylation state of GATC sequences, we perturbed the system in two independent ways. First, we increased the number of available GATC sites by transforming cells expressing MutH-Halo with the high-copy-number plasmid pUC19^46^. Each plasmid copy contains 19 GATC sites, and at an average copy number of ∼75 per cell at 37 °C, this corresponds to an additional ∼1400 GATC sites per cell relative to the wild-type chromosome. The fraction of immobile MutH trajectories increased significantly in the presence of the plasmid (Fig. 5a), demonstrating that MutH can bind these sites independently of chromosomal context.

**Fig. 5.**
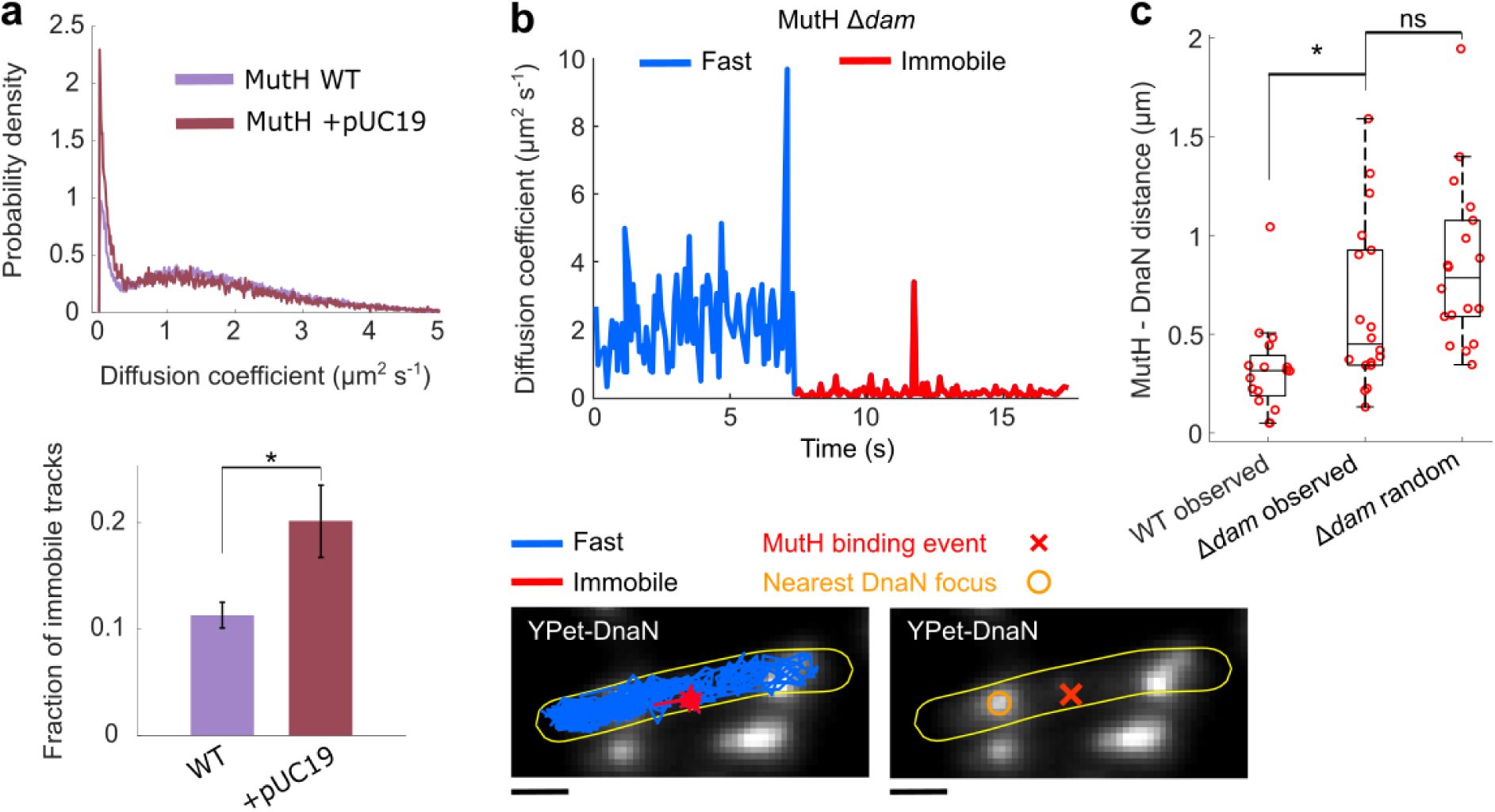
MutH binding is directed by hemimethylated GATC sites. **a,** Distribution of diffusion coefficients (top) and fraction of tracks classified as immobile (bottom) for MutH-Halo-TMR in wild-type cells and in cells transformed with plasmid pUC19. Distributions are pooled and bar plots show mean ± s.e.m from 3 independent experiments with pUC19, and 6 independent experiments for wild-type(*: P<0.05, ns: P>0.05). **b,** Top: Representative time-trace of diffusion coefficients and corresponding track of a single MutH-Halo-JFX650 molecule showing a binding event in a Δ*dam* cell. Bottom: Corresponding fluorescence snapshot of YPet-DnaN in the same cell. Localisations of the MutH binding event (red cross) and the nearest DnaN focus (orange circle) are shown. Scale bars, 1 µm. **c,** Box plots of the distances between MutH-Halo-JFX650 binding events and the nearest YPet-DnaN focus per cell. Observed are experimentally measured distances in wild-type and Δ*dam* cells. Random are distances when MutH binding sites were reassigned to random locations within the segmented cell area in Δ*dam* cells (* P<0.05, ns: P>0.05).

We next examined the role of GATC methylation in confining MutH binding to newly replicated DNA. To this end, we deleted the Dam methylase, such that all GATC sites across the chromosome remain unmethylated. MutH is known to bind and nick both unmethylated and hemimethylated GATC sites^47^, but not fully methylated sites^48^. In the absence of Dam, we therefore expected MutH binding to become distributed across unmethylated GATC sites throughout the entire chromosome. The duration of binding events of MutH in wild-type and Δ*dam* cells was similar (Fig. S4A). However, in Δ*dam* cells, MutH binding events were indeed located significantly further from DnaN foci than in wild-type cells (Fig. 5b,c), and the resulting distance distribution was indistinguishable from that of a randomised control (Fig. 5c). These results rule out replisome-mediated recruitment of MutH and instead show that methylation asymmetry at GATC sequences directs MutH binding to regions near replication forks in wild-type cells. Hemimethylation therefore not only solves the strand-discrimination problem, but also spatially guides MutH target search.

## Discussion

Efficient targeting of MutH to hemimethylated GATC sites is central to the function of the MMR pathway in *E. coli*. Incision of the newly synthesised strand must occur within minutes after passage of the replication fork, before Dam-mediated methylation erases the transient strand-discrimination signal. While the early steps of mismatch recognition by MutS and recruitment of MutL have been extensively characterised, the mechanisms that position and activate MutH remain less well defined. DNA incision by MutH constitutes the key committed step of the MMR pathway. Yet, this step is difficult to interrogate even under *in vitro* conditions with purified molecules because it is transient, spatially separated from the mismatch, and not readily synchronised. Live-cell single-molecule tracking can overcome these challenges, because the method enables direct observation of dynamic and short-lived processes in their native cellular context without population averaging or artificial synchronisation.

Previous *in vitro* studies have suggested that MutH may be delivered to the nearest hemimethylated GATC site by a sliding MutL or MutS–MutL complex following mismatch recognition^49^. Given that MutL serves as a central interaction hub within the MMR pathway, coordinating multiple downstream repair proteins^50^, it has seemed plausible that MutL also recruits MutH to its incision sites. However, these observations do not exclude an alternative scenario in which MutH binds hemimethylated GATC sites on its own and MutL functions primarily as an activator rather than a recruitment factor. Indeed, DNA incision by MutH can occur with similar efficiency whether the mismatch and GATC site are located on the same or on separate DNA molecules^51^, and MutH has intrinsic affinity for hemimethylated GATC sequences in the absence of mismatches or other MMR proteins with a kD below 1 µM^47^. Our results support a model in which MutH associates with hemimethylated GATC sites independently and does not require delivery by a MutS–MutL complex, consistent with activation through transient rather than long-lived ternary interactions. In this model, MutH trails the replication fork by repeatedly binding and dissociating from hemimethylated GATC sites on a timescale of seconds, irrespective of mismatch presence.

This mechanism likely offers several functional advantages. First, the search for hemimethylated GATC sites by MutH proceeds in parallel with mismatch detection by MutS, rather than requiring hierarchical recruitment. Pre-localisation of MutH near replication forks eliminates the need for long-range transport of a multiprotein complex over hundreds to thousands of base pairs, enabling rapid incision within the narrow temporal window before Dam-mediated methylation. Confining MutH binding to replication-coupled, hemimethylated regions also enhances specificity by restricting incision to sites where strand discrimination is unambiguous. Moreover, transient activation of pre-bound MutH by MutL minimises stoichiometric and organisational constraints, increasing robustness in the crowded intracellular environment. Finally, local enrichment of MutH near replication forks may increase the likelihood of multiple strand incision events around a mismatch, which have been shown to improve the efficiency of downstream DNA unwinding and degradation^52^.

The stoichiometry of MMR enzymes is consistent with a robust yet economical strategy. The number of MutH copies per cell is lower than the number of MutS or MutL dimers^53^. With approximately 100 MutH molecules per cell, an immobile fraction of ∼10% corresponds to only a few molecules associated with each replication fork at any given time. Such low occupancy is nevertheless sufficient to ensure rapid incision while limiting inappropriate cleavage elsewhere on the chromosome. Biochemical studies have shown that multiple MutH incision events can occur processively during a single repair reaction, increasing overall repair efficiency^52^. The presence of several MutH molecules in the vicinity of each replication fork would provide a simple means to support such processive incision activity *in vivo*. Considering the small number of MutH molecules, the high efficiency of mismatch repair is striking, especially when considering that many DNA-binding proteins fail to engage their targets even after reaching the correct chromosomal region, particularly when target sequences are short and sparsely distributed^54,55^. Our data provide an explanation: Frequent non-specific interaction of MutH with DNA facilitate an efficient target search^56^. Moreover, MutH switches from fast diffusion to a slow-search mode in the vicinity of replication forks. This slow-down is likely accompanied by increased DNA interaction, which raises the probability of productive encounters with hemimethylated GATC sites. In this framework, hemimethylation functions not only as a licensing signal for strand incision, but also as a spatial cue that directs MutH target search.

Together, our findings show that replication-coupled methylation asymmetry and spatially regulated search dynamics position MutH to execute strand incision with speed and precision in living cells. Outside γ-proteobacteria, a dedicated strand-incision factor such as MutH is absent and strand discrimination is achieved by alternative mechanisms^57,58^. In this light, *E. coli* may be more similar to other systems than previously appreciated, in that long-lived MutS– MutL–MutH ternary complexes are not a defining feature of the pathway.

**Fig. 6.**
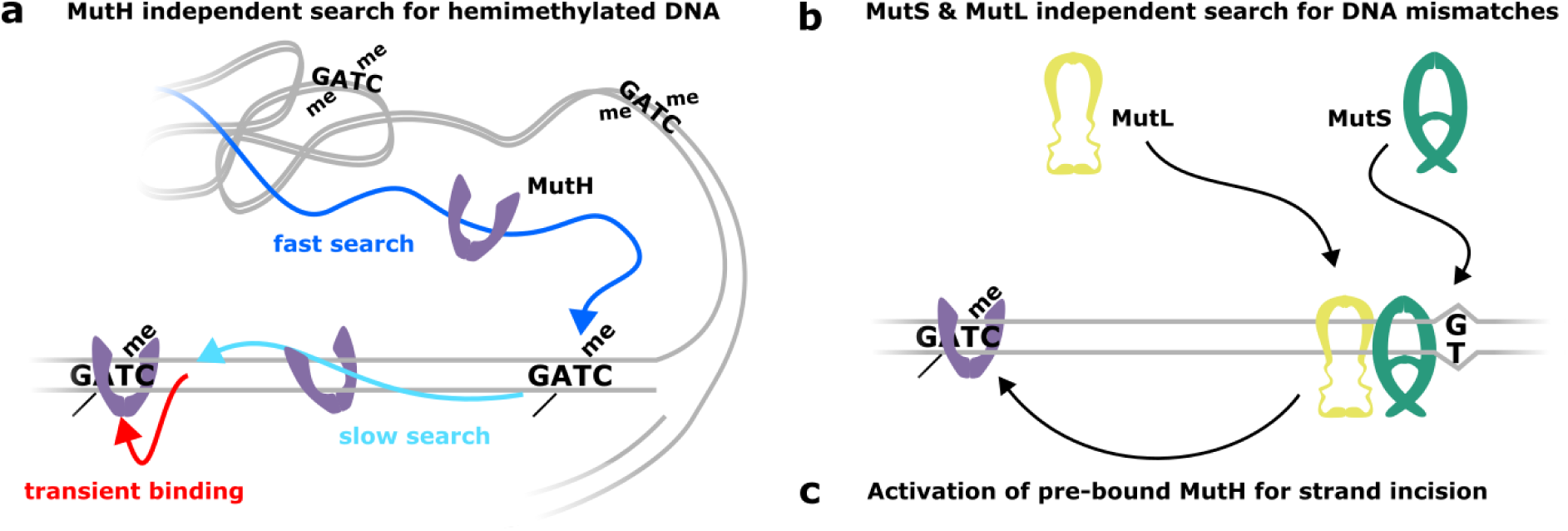
Parallel replication-coupled search positions MutH for strand incision in mismatch repair. **a,** MutH independently searches for hemimethylated GATC sites, switching from fast to slow search modes near DNA replication forks to guide target localisation. **b,** MutS locates DNA mismatches and recruits MutL, independently of MutH target search. **c,** Following mismatch recognition, MutS–MutL transiently activate pre-bound MutH to incise the unmethylated daughter strand.

## Acknowledgements

We thank all members of the Uphoff lab, past and present, for discussions. This research was funded by a Wellcome Trust Sir Henry Dale Fellowship (206159/Z/17/Z), and a Biotechnology and Biological Sciences Research Council grant (BB/ Y001567/1). We thank Joyce Lebbink, Titia Sixma, Peter Friedhoff, and all members of the RepState (DNA Repair State Machines) Marie Skłodowska-Curie Actions (MSCA) Doctoral Network for discussions.

## Methods

### Bacterial strain construction

All bacterial strains are derivatives of *E. coli* AB1157. We replaced the endogenous *mutS*, *mutL* or *mutH* genes with translational fusions to mYPet or the HaloTag^59^ using Lambda Red recombination^60^. Fusions included a flexible 11-amino acid linker between the protein C-terminus and the HaloTag and were followed by a kanamycin resistance cassette. Each fusion allele was moved into the wild-type AB1157 strain by P1 phage transduction, and the kanamycin resistance cassette was subsequently removed by expressing Flp recombinase from plasmid pCP20. The temperature-sensitive plasmid was transformed via heat shock transformation and maintained at 20 °C with selection for carbenicillin resistance. Shifting cells to 37 °C induced Flp recombinase and cured the plasmid. We selected for cells which had lost antibiotic resistance to both kanamycin and carbenicillin to produce strains SU358, SU359, and SU360 (for MutS-Halo, MutL-Halo, MutH-Halo), and SU171, SU172, and SU1518 for the corresponding mYPet fusions. Fusions were validated by colony PCR and through fluorescence imaging for successful labelling. Full mismatch repair functionality of the fusion proteins was confirmed by measuring the frequency of spontaneous rifampicin resistance mutations compared to wild-type and deletion mutant strains (see below).

To combine MutH-HaloTag fusion with targeted gene deletions, we used P1 phage transduction from donor strains JW2703 Δ*mutS:kan*, JW4128 Δ*mutL:kan*, and JW3964 Δ*hupA:kan* obtained from the KEIO collection, generating strains SU1620, SU413, and SU1746. We used the donor strain RR190 to create strain SU472 expressing MutH-Halo and *dnaN-mYpet:kan*. The Δ*dam:kan* allele from KEIO strain JW3350 was added to this strain via a second round of kanamycin resistance cassette removal and P1 phage transduction. The pUC19 plasmid was added to strain SU360 via heat shock transformation.

| Strain number: | Description: |
| --- | --- |
| SU000 | AB1157 wild-type (WT) |
| SU189 | AB1157 mutS-HaloTag:kan |
| **SU190** | AB1157 mutL-HaloTag:kan |
| **SU191** | AB1157 mutH-HaloTag:kan |
| **SU358** | AB1157 mutS-HaloTag (frt kan removed from SU189) |
| **SU359** | AB1157 mutL-HaloTag (frt kan removed from SU190) |
| **SU360** | AB1157 mutH-HaloTag (frt kan removed from SU191) |
| **SU171** | AB1157 ΔflhD, mKate2, mutS-mYpet:kan |
| **SU172** | AB1157 ΔflhD, mKate2, mutL-mYpet:kan |
| **SU1518** | AB1157 ΔflhD, mKate2, mutH-mYpet:kan |
| **SU515** | AB1157 ΔmutS::kan |
| **SU411** | AB1157 ΔmutL::kan |
| **SU412** | AB1157 ΔmutH::kan |
| **SU1620** | AB1157 mutH-HaloTag, ΔmutS::kan |
| **SU413** | AB1157 mutH-HaloTag, ΔmutL::kan |
| **SU1505** | AB1157 mutH-Halo (SU360) carrying pUC19 plasmid |
| **SU472** | AB1157 Ypet-dnaN:kan mutH-HaloTag |
| **SU545** | AB1157 Ypet-dnaN mutH-HaloTag (frt kan removed from SU472) |
| **SU1629** | AB1157 Ypet-dnaN, mutH-HaloTag, Δdam::kan |
| **SU1746** | AB1157 Ypet-dnaN, mutH-HaloTag, ΔhupA::kan |

### Mutation frequency assay

Rifampicin-resistance mutation frequency assays were used to assess the functionality of HaloTag- and mYPet-tagged MMR proteins. Mutations conferring rifampicin resistance (predominantly in the *rpoB* gene) provide a readout of spontaneous mutation frequency. Cells were streaked from glycerol stocks onto LB plates and grown overnight. Three single colonies per strain were inoculated into LB and grown overnight, then diluted 1/100 and grown to OD600 ≈ 0.8. 200 µl of cell suspension were plated onto LB agar containing rifampicin (100 µg ml⁻¹), and serial dilutions were plated onto LB agar without antibiotic to determine total viable cell counts. After overnight incubation, colonies were counted and mutation frequencies were calculated as the ratio of rifampicin-resistant colonies to total colony-forming units. Experiments were performed on three independent days with three biological replicates per condition (n = 9). Mutation frequencies of HaloTag- and mYPet-fusion strains were compared to wild-type *E. coli* AB1157 and corresponding gene deletion strains. Fusion strains exhibited mutation frequencies comparable to wild type and substantially lower than deletion strains (Fig. S.1), indicating that the tagged proteins retain mismatch repair activity.

### Bacterial culture

*E. coli* strains were stored at -70°C as frozen glycerol stocks, and streaked on LB agarose plates with appropriate antibiotic (25 μg/mL kanamycin for most strains in this study; 250 μg/mL carbenicillin for strain SU1505). For each experiment, a single colony was isolated and grown in LB media for ∼4 hours before being diluted 1/2000 in M9 minimal media (see below) and grown overnight. The following day, overnight cultures were diluted 1/50 and grown to early exponential phase for approximately 2 hours reaching OD600 ∼0.3 before labelling.

*E. coli* cultures were grown at 37°C in M9 minimal media for live-cell imaging. To make M9 media, the following were added to 100 mL of MilliQ water in autoclaved glass bottles: 500 µL 100 mM CaCl_2_, 1 mL 1M MgSO_4_, 100 mL 5x M9 minimal salts (Sigma-Aldrich), 10 mL 50x MEM amino acids (Gibco), 5 mL 10 mg/ml L-Proline, 50 µL 0.5% Thiamine, 5 mL 20 % Glucose; these were then topped up with MilliQ water to make a total volume of 500 mL M9 medium. The medium was filtered (Nalgene), and the stock stored at 4 °C. The volume of M9 media required for each experiment was decanted from the stock and warmed to room temperature prior to use, and the stock was remade weekly.

### Preparation of cells for single-molecule tracking

We labelled HaloTag fusion proteins using a protocol described previously^31^. HaloTag TMR ligand (Promega) was diluted from a 5 mM stock solution to a working concentration of 50 µM in MilliQ water. We used a 10x lower labelling concentration in experiments utilising HaloTag JFX650 ligand, with a 50 µM stock solution diluted to 5 µM working concentration in MilliQ water. 4 mL of bacterial culture was centrifuged and resuspended in 100 µL M9 media to form a concentrated suspension. 5 µL of the respective dye solutions were added and incubated with the cell suspension at room temperature for 30 minutes. Three washes were performed by repeated centrifugation and resuspension in 1 mL M9 to remove unbound dye. Cells were recovered for 30 minutes shaking in 2 mL M9 at 37 °C. A final wash with 1 mL M9 was performed following recovery, before cells were concentrated by centrifugation and spotted onto M9 agarose pads (see below). This procedure resulted in a density of 100-200 bacterial cells per field of view.

For experiments involving MMS or antibiotics, an additional incubation period was performed to treat cells after the final post-recovery wash. Washed cells were resuspended in 1 mL M9 supplemented with 0.01% MMS, 15 µg/mL chloramphenicol, or 50 µg/mL rifampicin (all supplied by Sigma-Aldrich) and shaken at 37 °C for 20 minutes (MMS) or 1 hour (chloramphenicol, rifampicin), respectively. Treated cell culture was concentrated by centrifugation and spotted onto M9 agarose pads also containing 0.01% MMS, 15 µg/mL chloramphenicol, or 50 µg/mL rifampicin to maintain constant treatment before and during imaging.

Agarose pads were made by adding Molecular Biology Agarose (Bio-Rad) to M9 minimal media (to 1% concentration) and gently heating in a microwave. Approximately 1 mL M9 agarose was then pipetted onto Borosilicate coverglass (#1.5 thickness, VWR International), and then sandwiched by a second coverglass on top, and left to solidify. Immediately before imaging, the top coverglass was removed, cells were spotted directly onto the agarose pad, and a new coverglass was placed on top of the cells. Coverglass were pre-cleaned using plasma etching (Plasma Etch PE-50).

### Single-molecule tracking instrumentation

For all single-molecule tracking experiments, we used a custom inverted optical microscope assembled on a Rapid Automated Modular Microscope body (ASI Applied Scientific Instrumentation) consisting of: a multilaser engine (iChrome MLE, Toptica Photonics) with fibre-coupled 405, 488, 561, 640 nm lasers, EMCCD camera (iXon Ultra 897 with 300x gain, Andor), Plan Apo λ 100x / 1.45 oil objective (Nikon), motorized stage for sample mounting and XY translation with piezo insert for focus control (ASI), quadband dichroic mirror (ZT405/488/561/640rpc, Chroma), emission filter wheel (Chroma, 425-475 nm bandpass, 500-550 nm bandpass, 575 nm longpass, 655 nm longpass), LED condenser for transmitted light illumination (Nikon). Optomechanical components were controlled by a TG-1000 unit (ASI) via Micromanager^61^ and Solis software (Andor)., we used a pair of lenses (focal lengths 180 mm and 250 mm) to position the laser focus onto the back focal plane of the objective. The laser fibre output and lens pair were mounted on a translatable stage with motorized actuator (MB1530F/M, z812, KDC101, Thorlabs) to shift the angle of the illumination beam for Highly Inclined and Laminated Optical sheet (HILO) illumination. We cropped the camera field of view to 300 x 512 pixels, at a pixel resolution of 107 nm generated by a 300 mm tube lens.

### Single-molecule tracking data acquisition

Cells were first illuminated with an LED condenser to capture a transmitted light image (exposure time 30 ms), which aids cell segmentation during the analysis of single-molecule trajectories. For single-molecule tracking acquisitions, we used an exposure time of 15 ms/frame for MutH-Halo and MutL-Halo, and 30 ms/frame for MutS-Halo throughout this study. The camera readout process for the field of view size adds 0.475 ms to the total frame intervals.

For single-molecule tracking using TMR ligand, we used continuous excitation with a 561 nm laser using a power density at the sample plane of approximately 1 W cm^-2^. At each field of view, cells were initially exposed for a brief period, during which TMR fluorophores deactivate and the density of fluorescent spots reduces to sparse-emitter conditions of less than one spot per cell per frame. Following this, we recorded a movie of 10000 frames per field of view for MutH-Halo and MutL-Halo, or 5000 frames per field of view for MutS-Halo. Typically, 6-8 fields of view were imaged in this manner for each sample, providing data for >1000 cells per session. At least 3 independent imaging sessions with cultures grown from separate colonies were performed for each experimental condition presented in this work.

To measure long trajectories of single MutH-Halo molecules labelled with the photostable fluorophore JFX650, we recorded movies of ∼3000 frames (or until all fluorescent molecules photobleached in a field of view) at 15 ms/frame immediately from the moment of 640 nm laser activation (power density approximately 1 W cm^-2^) without prior illumination of cells. The lower concentration JFX650 used for labelling ensured that sparse-emitter conditions were achieved from the start of each movie. We imaged at least 12 fields of view per session.

In experiments involving the YPet-DnaN fluorescent marker for replication loci, we illuminated cells with the 488 nm laser at a power density below 1 W cm^-2^ between capturing the transmitted light image and recording the single-molecule tracking movie. During the 488 nm illumination, we acquired a short movie of approximately 1000 frames with 15 ms exposure time, whilst simultaneously manually sweeping the sample focus to record a tomogram of the cell volume. Average intensity projections of the movie frames then formed an image of YPet-DnaN foci locations.

### Single-molecule tracking analysis

Single-molecule imaging data were processed and analysed using MATLAB-based scripts. The transmitted light image for each field of view was used to segment the outlines of *E. coli* cells with microbeTracker. We excluded any dead cells that had taken up large quantities of fluorophore. Next, single spots are localized in each frame of the single-molecule tracking movie by setting a minimum intensity threshold for point spread function detection and determining the centroid position using a phasor localization algorithm^62^. Detected spots are linked together to form single-molecule tracks if the localizations are within the same cell volume and the distance between localizations in subsequent frames is less than 8 pixels. A memory parameter of 1-frame accommodated missed localisations in single frames of a track. Single-molecule trajectories were then split into sub-trajectories each of 5 frames to calculate the mean squared displacement (MSD) and diffusion coefficient, using 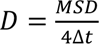 with a frame interval of Δt. Splitting trajectories in this manner reduces sampling biases of molecules moving at different speeds^30^.

For JFX650 tracks, sub-trajectories were concatenated to produce plots of D vs. time. For TMR tracks, all diffusion coefficient measurements from multiple fields of view per independent repeat were accumulated into a single histogram. A mixture of probability density functions was fit to the histogram to estimate the relative fractions of molecules in different diffusion states. For each state, the distribution of diffusion coefficients was modelled by a Gamma distribution:

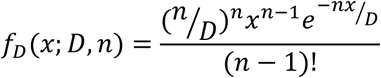

where *x* is the diffusion coefficient of an individual molecule, *D* is the average diffusion coefficient, and *n* is the number of steps the mean-square displacement (MSD) is being calculated over (here 4 steps). Least-squares fitting of a sum of two Gamma distributions (immobile, mobile) or three Gamma distributions (immobile, slow-diffusing, fast-diffusing) provides estimates for the average diffusion coefficient and relative proportions of each state^63^. We constrained the average diffusion coefficient of the immobile population and kept the diffusion coefficients of the mobile populations unconstrained. The initial fit parameters and constrained values were obtained by analysing the tracking data separately using vbSPT^64^.

## Notes

### Competing Interest Statement

The authors have declared no competing interest.

