## Supplementary information for "Replication-coupled search positions MutH for strand incision in DNA mismatch repair"

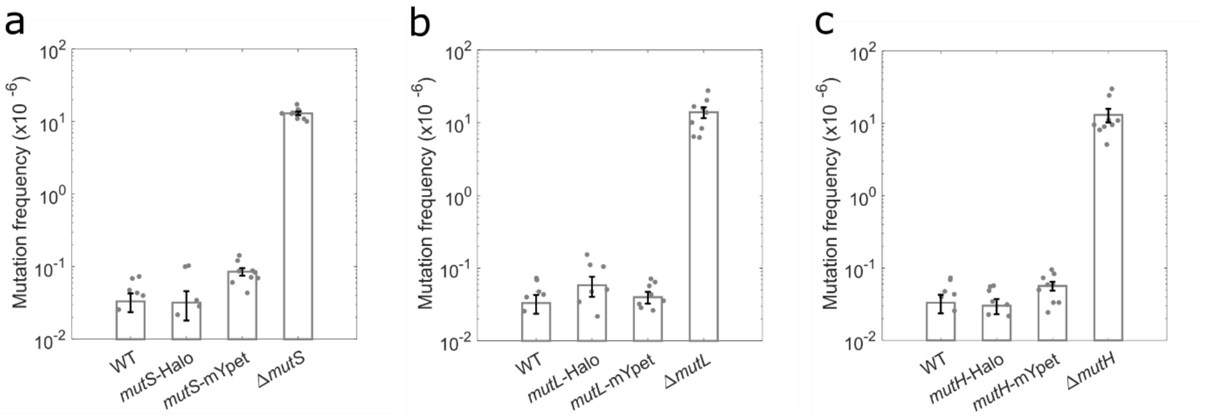


**Extended Data Fig. 1. Functionality of MMR fusion constructs.** Spontaneous mutation frequencies calculated from the ratio of rifampicin-resistant colonies to total colony-forming units for *E.coli* AB1157 WT and strains expressing HaloTag, mYPet, or deletions of *mutS* (a), *mutL* (b) and *mutH* (c) enzymes. Bar heights and error bars represent mean ± SEM of 9 independent cultures (dots).

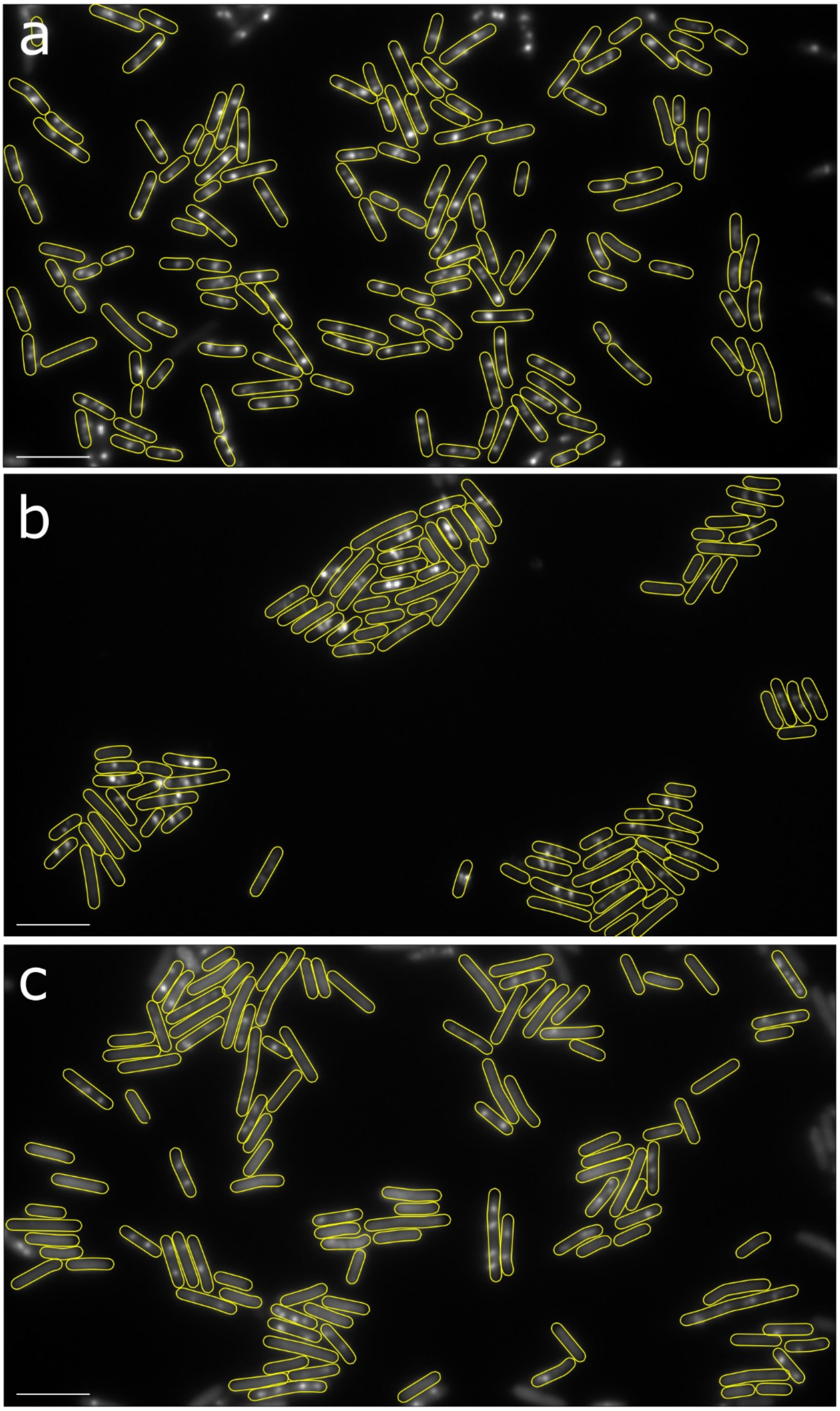


**Extended Data Fig. 2. Snapshots of YPet-DnaN foci**, indicating locations of active DNA replication, in untreated cells (a), and cells treated with 15 μg/mL chloramphenicol (b) or 50 μg/mL rifampicin (c) for 1 hour. Segmented cell outlines shown in yellow. Scale bars: 5 μm.

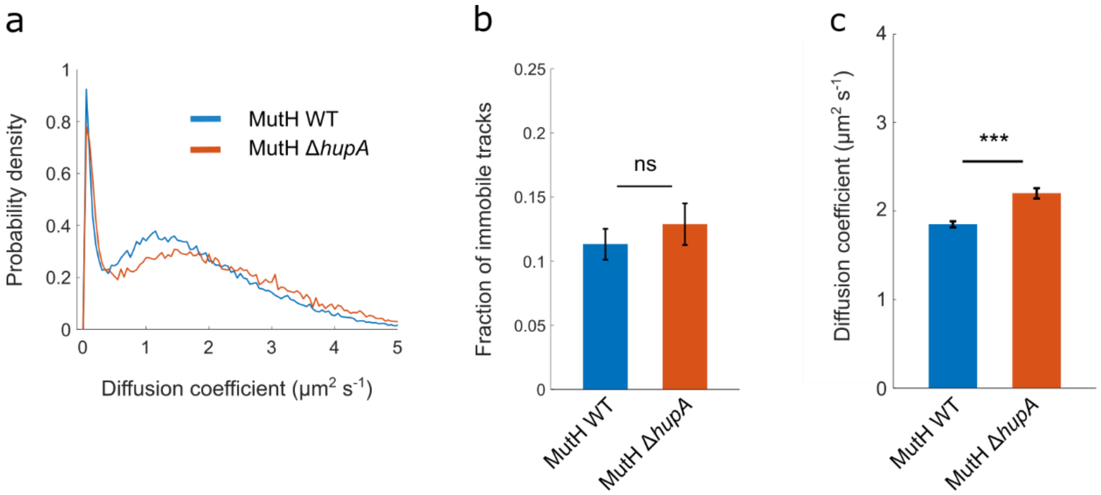


**Extended Data Fig. 3. Increased diffusion coefficient of MutH in the absence of HU.** (a) Distribution of diffusion coefficients for MutH-Halo-TMR in wild-type cells (blue) and in cells with Δ*hupA* gene deletion (red). Data are pooled from 6 independent experiments for wild-type and 3 independent experiments for Δ*hupA*. (b) Fraction of MutH-Halo-TMR tracks classified as immobile in wild-type and Δ*hupA* cells (ns: P>0.05). (c) Mean diffusion coefficient of diffusing MutH-Halo-TMR molecules in wild-type and Δ*hupA* cells (***: P<0.0005).

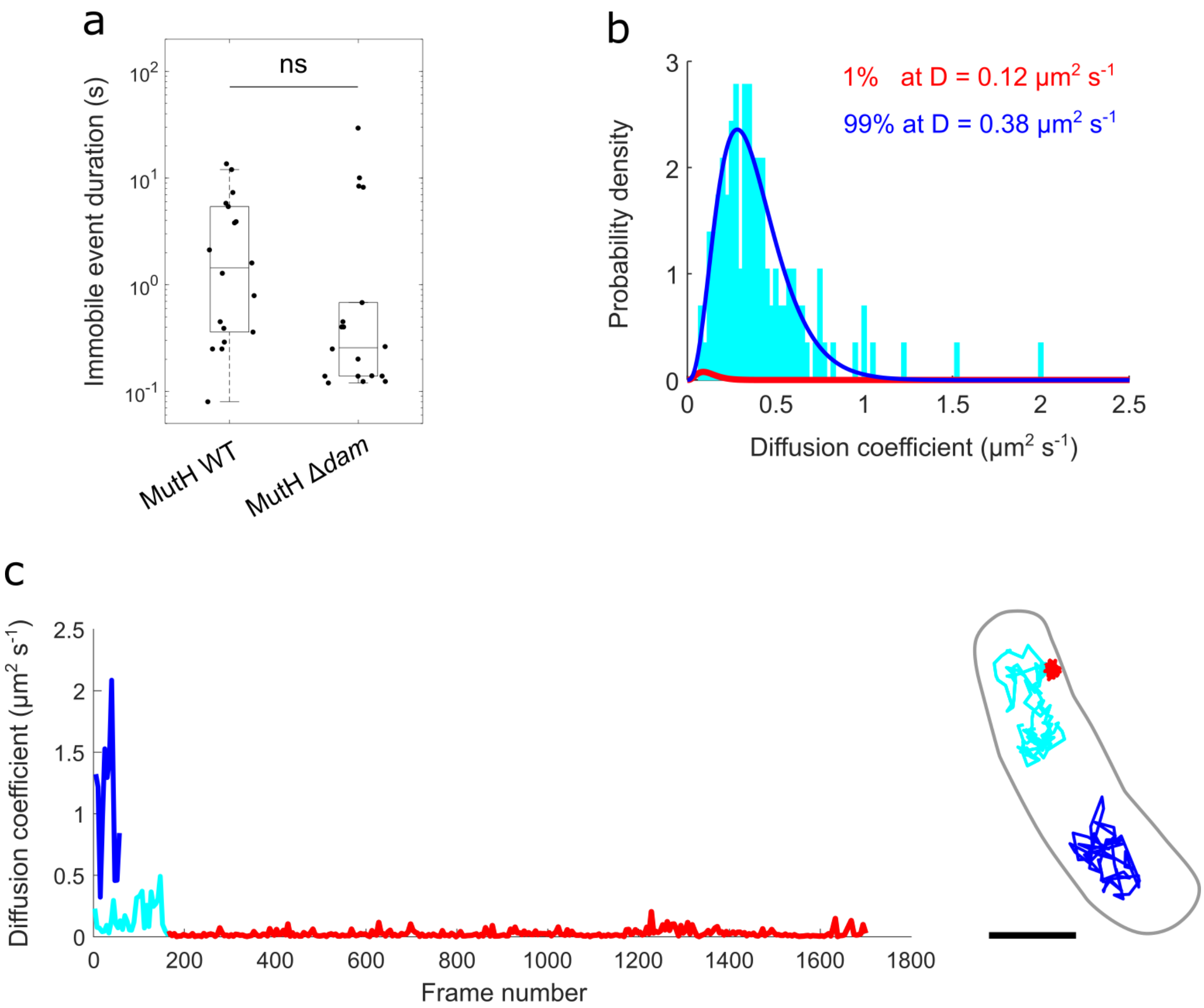


**Extended Data Fig. 4. Analysis of MutH-Halo diffusion characteristics under different conditions.** (a) Durations of immobile states of MutH-Halo-JFX650 in wild-type (WT) and Δ*dam* strains. Box plot show individual immobile events with mean, upper and lower quartiles (ns: P>0.05). (b) Validation of the two-state model used to estimate the fraction of immobile MutH molecules. Long-lived MutH-Halo-JFX650 trajectories revealed a previously unresolved slow-diffusing state in addition to the immobile and fast-diffusing states. Fitting the diffusion coefficient distribution of a representative slow-diffusing trajectory showed that this state makes a negligible contribution to the immobile population, validating the two-state analysis used to quantify MutH binding. (c) Immobile (red), slow-diffusing (cyan), and fast-diffusing (blue) states of MutH still exist when MutL is deleted. Representative time-trace of diffusion coefficients and corresponding track of two single MutH-Halo-JFX650 molecules in a cell with Δ*mutL* deletion. Scale bar: 1 μm.
